# Short-term effects of sound localization training in virtual reality

**DOI:** 10.1101/207753

**Authors:** Mark A. Steadman, Chungeun Kim, Jean-Hugues Lestang, Dan F. M. Goodman, Lorenzo Picinali

## Abstract

Head-related transfer functions (HRTFs) capture the direction-dependant way that sound interacts with the head and torso. In virtual audio systems, which aim to emulate these effects, non-individualized, generic HRTFs are typically used leading to an inaccurate perception of virtual sound location. Training has the potential to exploit the brain’s ability to adapt to these unfamiliar cues. In this study, three virtual sound localization training paradigms were evaluated; one provided simple visual positional confirmation of sound source location, a second introduced game design elements (“gamification”) and a final version additionally utilized head-tracking to provide listeners with experience of relative sound source motion (“active listening”). The results demonstrate a significant effect of training after a small number of short (12-minute) training sessions, which is retained across multiple days. Gamification alone had no significant effect on the efficacy of the training, but active listening resulted in a significantly greater improvements in localization accuracy. In general, improvements in virtual sound localization following training generalized to a second set of non-individualized HRTFs, although some HRTF-specific changes were observed in polar angle judgement for the active listening group. The implications of this on the putative mechanisms of the adaptation process are discussed.

## Introduction

Sounds interact with the head and torso in a direction-dependant way. For example, sounds sources located to the side will reach the contralateral ear after a longer delay relative to the ipsilateral ear, and with lower intensity. Furthermore, physical interactions with the head and pinnae, the external parts of the ear, introduce spectral peaks and notches, which can be used to judge whether a sound source is above, below or behind the listener. This direction-dependant filtering is described by Head-Related Transfer Functions (HRTFs). Virtual audio systems are based on the premise that, if the HRTFs for a given listener can be effectively estimated, any monoaural sound can be processed in such a way that, when presented over headphones it is perceived as if it emanates from any position in 3D space^1^.

Because of individual differences in the size and shape of the head and pinnae, HRTFs vary from one listener to another. It follows that an ideal virtual audio system would make use of individualized HRTFs. This is problematic for virtual audio systems designed for use in consumer or clinical applications, because the equipment required to measure HRTFs is typically bulky and costly. Some work has been done on estimating HRTFs from readily accessible anthropometric information; for example, measurements of the pinnae and head^2, 3^ or even photographs^4, 5^. However, such approaches necessitate the use of simplified morphological models, the limitations of which are unclear. The most accurate estimations of HRTFs typically involve the use of specialized equipment, ranging from rotating listening platforms to spherical loudspeaker arrays and robotic armatures (for a brief overview see Katz & Begault, 2006^6^) along with miniature, accurate microphones that can be placed inside the ear. For this reason, consumer-oriented systems typically use generic HRTFs measured from a small sample of listeners, or artificial anthropometric models such as the KEMAR head and torso^7^.

It is generally thought that the differences between individualized HRTFs and these generic ones have a detrimental effect on the accuracy and realism of virtual sound perception. It has been noted, for example, that listeners are able to localize virtual sounds that have been spatialized using individualized HRTFs with a similar accuracy to free field listening, albeit with somewhat poorer elevation judgments and increased front-back confusions^1, 8^. These errors are exacerbated by the use of non-individualized HRTFs^9, 10^. Furthermore, it has been suggested that non-individualized HRTFs result in an auditory perception with reduced “presence”^11^. It would seem that the efficacy of virtual audio systems utilising generic HRTFs is limited by the perceptual similarity between these HRTFs and the listener’s own. Efforts have been made to “match” listeners to optimal HRTFs from a database using subjective methods^12–14^. Whilst this is a promising, efficient approach, it does not take advantage of the brain’s ability to adapt to changes in sensory input. There is increasing evidence that the adult brain is more adaptable than classically thought^15^. For example, it has been demonstrated that this adaptability (or plasticity) can lead to a decrease in localization error over time when a listener’s normal cues for sound location are disrupted by physically altering the shape of the ear using molds^16–18^. However, this process typically takes place over the course of days or weeks (for a review see Mendonça *et al.*^19^).

Such timescales are likely to be impractical for a consumer-oriented or clinical applications, where rapid optimization is generally desirable. The possibility of accelerating the process of adapting to “new ears” has therefore received some attention. Encouragingly, several studies have demonstrated that training through positional feedback (for example, indication of virtual sound source location using visual or somatosensory cues) has the potential to achieve adaptation over timescales of the order of a few hours or even minutes^20–24^. Whilst it seems clear that explicit training can result in better outcomes in virtual audio, whether measured by localization accuracy or perceived externalization, improvements over short timescales are typically small and highly variable.

It is possible that such training paradigms could be further optimised. One promising avenue of investigation to that end is the use of “gamification”, whereby design elements traditionally used in gaming are employed in a non-gaming context. Not only is gameplay engaging, having the potential to improve attention to a perceptual learning task, but it also leads to the release of reward signals such as dopamine^25^, which in turn have been purported to have an enhancing effect on perceptual learning through the promotion of synaptic plasticity in sensory processing nuclei^26^. The efficacy of video games to enhance various aspects of perceptual learning has been explored in the visual domain^27–30^ and, more recently, in the auditory domain^31–34^. However, to what extent gamification can accelerate virtual sound localization training relative to a more traditional approach is unknown.

Changes in virtual sound localization performance following training are typically described as “HRTF adaptation”^19, 23, 35^. However, the mechanisms underlying this adaptation are unclear. One possibility is that this adaptation reflects a process of learning a new, internal representation of the unfamiliar HRTFs in parallel to the listener’s own^16^. In this case, one would expect any changes in localization performance to be specific to the HRTFs used during the adaptation or training period. A second possibility is that the process may involve cue reweighting, whereby the listener learns to prioritise cues that remain robust despite perceptual differences between their own HRTFs and the generic set. If this is the case, listeners would be likely to prioritise cues that generalise to several generic HRTF sets. A cue reweighting mechanism has been reported in adult listeners in a sound localization study utilizing unilateral ear plugs^36^. Understanding the mechanism of HRTF adaptation could have implications for virtual audio system design and may be of interest in the field of auditory perceptual learning more generally.

This study will address the following questions. Firstly, we will examine whether virtual sound localization training using visual positional feedback can improve localization accuracy in virtual reality. We focus on the short-term effects of training (a total of 108 minutes, spaced over three days), since the study was motivated by considerations of potential consumer or clinical applications of virtual audio. Secondly, we investigate the efficacy of several training paradigm variants to effect these improvements. Specifically, we examine whether a “gamified” training task leads to significantly improved short-term training effects relative to a similar, non-gamified one. In addition, we also address the question of whether “active listening” (moving the head relative to a fixed virtual sound source) is an important factor in adapting to “new ears”. We hypothesize that both gamification and active listening should lead to improvements in training efficacy relative to a standard, non-gamified paradigm. We also examine the nature of any improvements by examining changes in lateral and polar angle judgements, as well as the changes in rates of front-back confusions and response biases.

Previous studies have compared improvements due to training for listeners using their own, individualized HRTFs with those using non-individualized HRTFs^23, 37^. Here, we investigate whether any learning effects, as a result of training, transfer to a second set of non-individualised HRTFs, for which the listeners received no positional feedback during training. We then discuss the possible implications of our findings in terms of the putative mechanisms underlying “HRTF adaptation”. Finally, considering that measurement of virtual sound localization accuracy often relies on the development of highly specialised systems that integrate sound presentation and head tracking hardware, we demonstrate that both measurement and training can be implemented using a virtual audio system comprising readily available consumer electronics.

## Results

In total, 36 participants were recruited for this study. These were divided into four groups, three of which underwent sound localization training; “non-gamified” (*n* = 9), “gamified” (*n* = 7), or “active-gamified” (*n* = 11). The final group acted as a control and did not undergo training (*n* = 9).

### Effects of virtual sound localization training

Virtual sound localization errors were measured before and after training to accurately localize sounds spatialized using non-individualized HRTFs presented over headphones. During testing, participants were presented with a spatialized stimulus after which they were required to indicate the perceived direction of the virtual sound by orienting towards it and pressing a button to indicate their response. This orientation was measured using embedded sensors in a smartphone-based, head mounted display. Between testing blocks, participants underwent virtual sound localization training, during which they were provided with visual positional feedback, indicating the true sound source location after each response. There was a total of nine, 12-minute training blocks split over three days. Additional testing blocks were carried out at the beginning and end of each day, and between every training block on the first day in order to capture the dynamics of any very rapid changes in localization accuracy. This section presents the changes that occurred over the entire course of training. The timescale of learning is addressed explicitly in a subsequent section.

A total of 36 participants were randomly assigned to one of four groups; a control group (*n* = 9) that did not complete any training blocks, and three training groups. Each training group utilised a different variant of the training software, referred to as “non-gamified” (*n* = 9), “gamified” (*n* = 7), and “active-gamified” (*n* = 11). The distributions of localization errors pooled across all participants within each group, as measured by the angle between the target and response orientations (hereafter referred to as the spherical angle error), are shown in the top row of Fig. 1. Within all groups and testing blocks, the errors tend to be skewed, with the majority of errors having a magnitude of <90°, with some extending to almost 180°(the maximum). For this reason, per-participant median errors are used as the dependent variable in subsequent statistical analyses, since this provides a better description of the central tendency of these distributions. The bottom row of Fig. 1 shows the localization error as a function of target azimuth before and after training. In all cases, the largest errors were observed when the virtual sound sources were located directly behind the participants. For example, the “gamified” participant group exhibited localization errors of over 90° when target sounds had an azimuthal angle close to 180° (directly behind). Although the errors are substantially reduced following training in all groups except the control, the largest errors still tend to occur for targets in this region.

**Figure 1.**
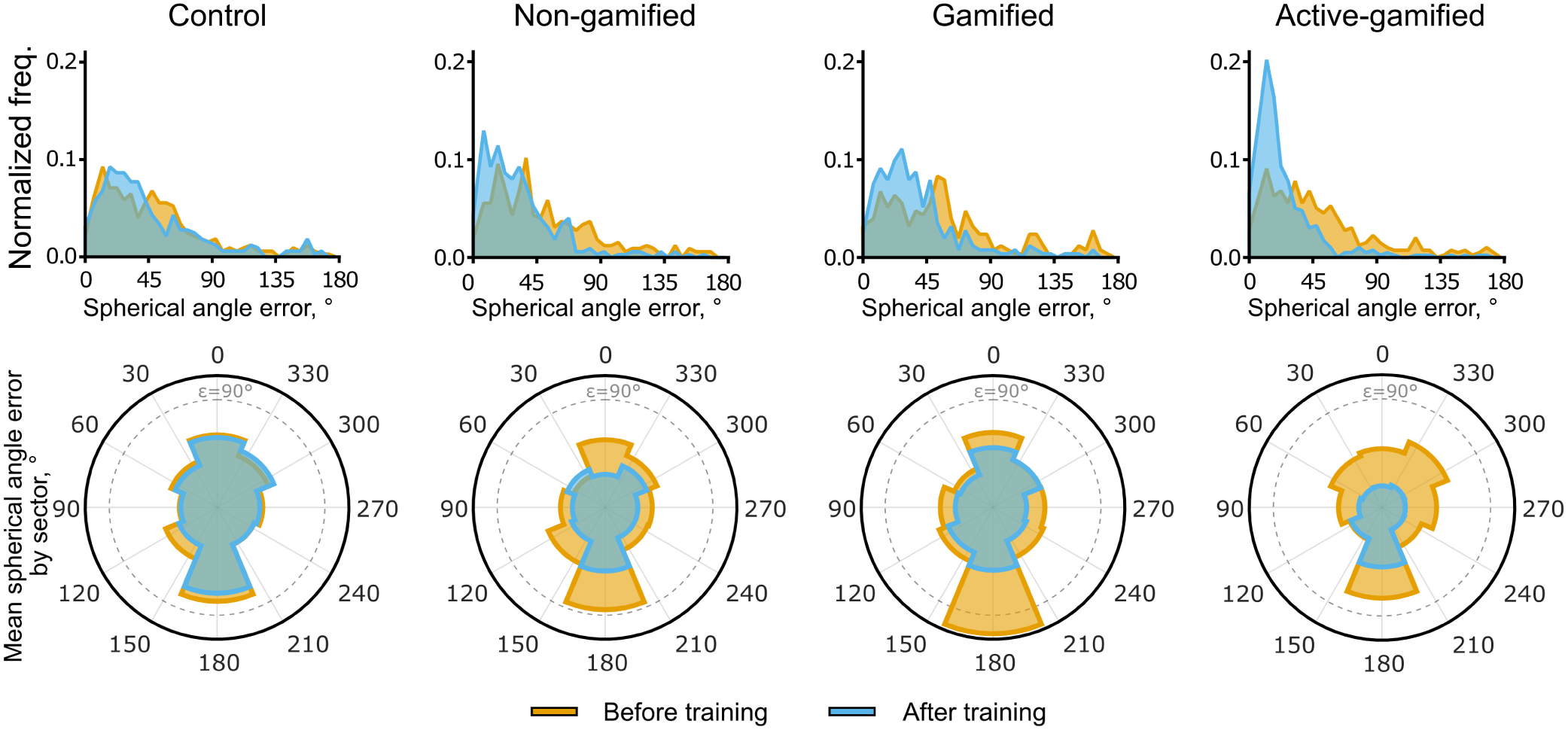
(**top row**) distribution of localization errors pooled across all participants within each group before training (orange) and after completing a total of nine, 12-minute training sessions across three days (or following a matching testing schedule without training for the control group). (**bottom row**) polar histograms of average localization error grouped by target azimuth into eight sectors both before (orange) and after training (blue). The dashed lines constitute the scale bars of the histograms and correspond to a mean localization error of 90°.

Localization errors before and after training, separated by training variant, are summarized in Fig. 2a. Separate paired t-tests confirmed that each training paradigm resulted in statistically significant reductions in localization error, while there was no significant reduction for the control group (*t*_*control*_ (8) = 1.72, *p* = 0.124; *t*_*non*−*gami f ied*_ (8) = 4.94, *p* < 0.0011; *t*_*gami f ied*_ (6) = 6.33, *p* < 0.0115; *t*_*active*−*gami f ied*_ (10) = 6.33, *p* < 0.001). In order to directly compare the training types, a one-way ANCOVA was conducted to determine whether there was a statistically significant difference between the final localization errors between groups, whilst controlling for any differences in initial localization error. This indicated a significant effect of participant group, *F*(3, 31) = 9.19, *p* < 0.001. Tukey *post hoc* tests revealed that the adjusted final localization error was lower for active-gamified group compared to all other groups, and that this difference was statistically significant when compared to the gamified and control groups, *p* < 0.001 and *p* = 0.024, respectively. These *post hoc* tests also indicated that the active-gamified and non-gamified groups had significantly lower final localization errors than the control group after controlling for initial performance (*p* < 0.001 and *p* = 0.05). The adjusted final localization error for gamified training group was also lower than that of the control group (Δ*µ* = 5.560°), but this difference was not statistically significant, *p* = 0.940.

**Figure 2.**
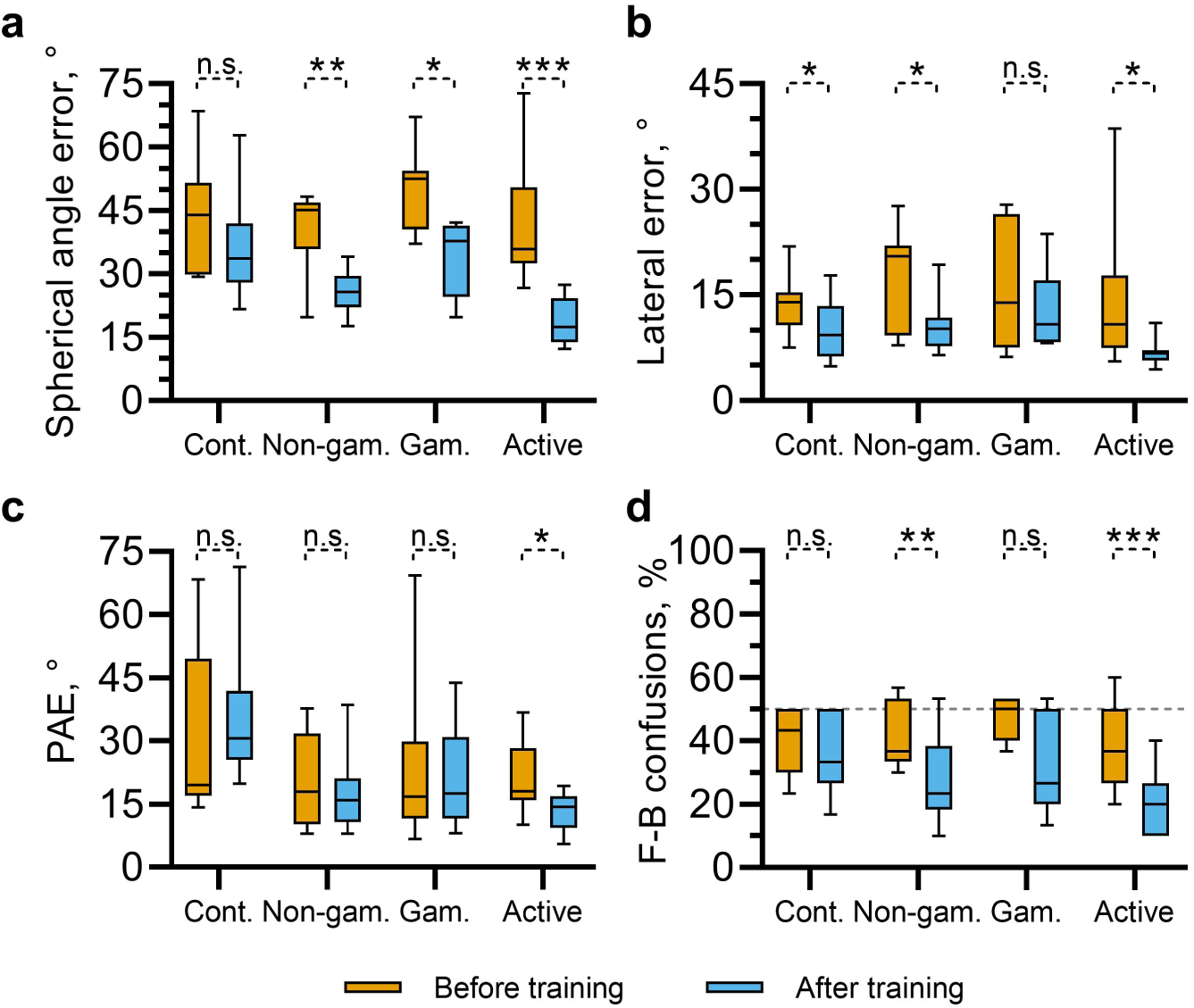
Distributions of localization errors, divided by participant group, during the initial (before training, orange), and final testing block (after training, blue). Shown are the spherical angle errors (**a**), lateral angle errors (**b**), polar angle errors (PAE; **c**) and the rates of front-back confusions (**d**). For all angle errors, the madian value was calculated for each participant in each testing block. Significance indicators show results of separate, paired t-tests between each initial and final measures for each participant group.

As described above, participants in the active-gamified group had the lowest adjusted final spherical angle errors. One factor that may contribute to this is the fact that target stimuli were played continuously throughout each trial in this training variant, while they were played only once whilst participants remained in a fixed position in the other variants. Furthermore, since participants engaged with training for a fixed duration, rather than a fixed number of trials, it was likely that there could be differences in the number of trials each participant completed. Indeed, a one-way ANOVA confirmed that the total number of stimulus presentations significantly differed between the groups that underwent training, *F*(2, 24) = 214.0, *p* < 0.001. However, there was no significant correlation between the total number of stimuli heard by each participant throughout training (across all groups) and the total reduction in spherical angle error from the initial to the final evaluation, *r* = −0.236, *p* = 0.235. This suggests that factors other than the total number of times each stimulus was heard are needed to account for the observed differences between training paradigms.

Improvements in localization accuracy as a result of training could be driven by adaptation to non-individualised timing or level differences between the ears, which serve as dominant cues for left-right position. They could also be driven by adaptation to novel spectral cues, such as the position of spectral notches, which vary dependant on source elevation and can facilitate resolution of front-back ambiguity. In order to investigate the relative contributions of these factors to the overall reduction in localization error, we used the auditory-inspired interaural polar coordinate system^8^ to define errors in terms of their lateral, polar and front-back components (see). A similar approach to that described above was used to analyse the effect of training on lateral error, polar angle error (PAE) and front-back (F-B) confusions separately. Separate t-tests were carried out for each measure and each participant group, the results of which are indicated in Figs. 2b-d.

For lateral angle errors (Fig. 2b), the separate, paired t-tests indicated a significant reduction in error between the initial and final testing blocks for participants using the non-gamified variant, *t*_*non*−*gami f ied*_ (8) = 2.441, *p* = 0.0018, and the active-gamified variant, *t*_*active*−*gami f ied*_ (10) = 2.69, *p* = 0.023. There was also a significant reduction in lateral error for participants in the control group, who did not undergo training, *t*_*control*_ (8) = 2.44, *p* = 0.040. Participants in the gamified group did not show a significant reduction, *t*_*gami f ied*_ (6) = 0.843, *p* = 0.432. A one-way ANCOVA indicated that there was a significant effect of participant group on lateral error after training, whilst accounting for differences in initial lateralization error, *F*(3, 31) = 3.385, *p* = 0.030. Tukey *post hoc* tests indicated that this effect was primarily driven by a significant difference in adjusted final lateral angle errors between the gamified and active-gamified groups, Δ*µ*_*ad j*_ = 5.69°, *p* = 0.024. Other pairwise comparisons were not significant.

For polar angle errors (PAE; Fig. 2c), separate paired t-tests indicated significant reduction in error between the initial and final testing blocks only for participants using the active-gamified variant, *t*_*active−gami f ied*_ (10) = 3.12, *p* = 0.011. The control, non-gamified and gamified groups showed no significant differences, *t*_*control*_ (8) = −1.955, *p* = 0.086; *t*_*non−gami f ied*_ (8) = −0.050, *p* = 0.962; *t*_*gami f ied*_ (6) = −1.313, *p* = 0.237. A one-way ANCOVA indicated a significant effect of group on PAE after training, whilst accounting for differences in initial PAE *F*(3, 31) = 5.761, *p* = 0.003. Tukey *post hoc* tests indicated that differences between each of the trained groups were not significant. However, the non-gamified and active-gamified groups differed significantly from the control group, *p* = 0.028, *p* = 0.002, respectively. The same tests indicated that the final PAE did not differ significantly between the gamified and control groups after taking initial performance into account, *p* = 0.227.

For front-back confusions (F-B confusions; Fig. 2c), separate paired t-tests indicated a significant reduction in error between the initial and final testing blocks for participants using the non-gamified and active-gamified training variants, *t*_*non−gami f ied*_ (8) = 4.296, *p* = 0.003; *t*_*active−gami f ied*_ (10) = 6.056, *p* < 0.001. Differences between the rate of front-back confusions from the initial to final testing block were not statistically significant for the group using the gamified training variant, *t*_*gami f ied*_ (6) = 2.121, *p* = 0.078, or the control group, *t*_*control*_ (8) = 0.973, *p* = 0.359. Despite these differences, a one-way ANCOVA indicated no significant effect of group on the rates of front-back confusions in the final testing block whilst accounting for those in the initial testing block, *F*(3, 31) = 2.411, *p* = 0.086.

In summary, all groups undergoing training showed lower localization errors on average following training than the control group (who only took part in testing blocks), after accounting for initial localization performance. This was most notable in the spherical angle error, which encompasses lateralization judgements, elevation judgements and front-back confusions in a single measure. Changes in PAE and front-back confusion rates yielded a similar pattern of results, although the variance within each group coupled with relatively small effects meant that these changes were not statistically significant in several cases. However, active listening appears to play an important role in the efficacy of training, since participants in this group robustly showed improvements in all aspects of localization judgements, whereas the other groups did not. Differences in the total number of stimulus presentations throughout training do not appear to wholly account for this.

### Changes in response bias

It was hypothesized that response biases might account for some of the observed changes in localization accuracy. In order to examine any systematic response biases, the signed lateral angle errors, signed PAEs and signed front-back confusions were examined. For signed lateral errors, a positive value would indicate a tendency to respond more laterally to the target angle and a negative value indicates a tendency to respond more medially. For signed elevation errors, a positive value indicates a tendency to give responses above the target and a negative value indicates a tendency to respond lower than the target. These metrics were only calculated for responses in the correct front-back hemisphere. Finally, for signed front-back (F-B) confusions, a positive value indicates a tendency to perceive the target in the front hemisphere when it was behind and a negative value indicates a tendency to perceive the target behind when it was in front.

Fig. 3a shows the signed lateral error. The mean signed lateral errors before training were above zero for all groups (*µ*_*control*_ = 10.7°, *µ*_*non−gami f ied*_ = 8.45°, *µ*_*gami f ied*_ = 10.9°, *µ*_*active−gami f ied*_ = 7.46°), indicating a tendency to respond more laterally than the target. On average, the magnitude of this error was reduced after training. Separate, paired t-tests indicated that this change was statistically significant for the gamified and active-gamified groups; *t*_*gami f ied*_ (6) = 2.85, *p* = 0.029, *t*_*active−gami f ied*_ (10) = 2.62, *p* = 0.026. Signed elevation errors are shown in Fig. 3b. Here, all groups showed a strong tendency to perceive sound source elevation lower than the target during the initial testing block (before training; *µ*_*control*_ = −20.3°,*µ*_*non−gami f ied*_ = −11.3°, *µ*_*gami f ied*_ = −16.8°, *µ*_*active−gami f ied*_ = −11.3°). This tendency was reduced following training for all groups undergoing training, leading to the significant changes in the signed elevation error for the non-gamified, gamified and active-gamified groups; *t*_*non−gami f ied*_ (8) = −3.24, *p* = −0.012, *t*_*gami f ied*_ (6) = −2.655, *p* = 0.038, *t*_*active−gami f ied*_ (10) = −4.071, *p* −0.002. There appeared to be no strong tendency towards front to back rather than back to front confusions (Fig. 3c), although the active-gamified group did show a reversal from a small front to back preference (*µ* = −0.049) to a small back to front preference (*µ* = 0.124), and a paired t-test indicated that this change was statistically significant; *t*(10) = −2.738, *p* = 0.021.

**Figure 3.**
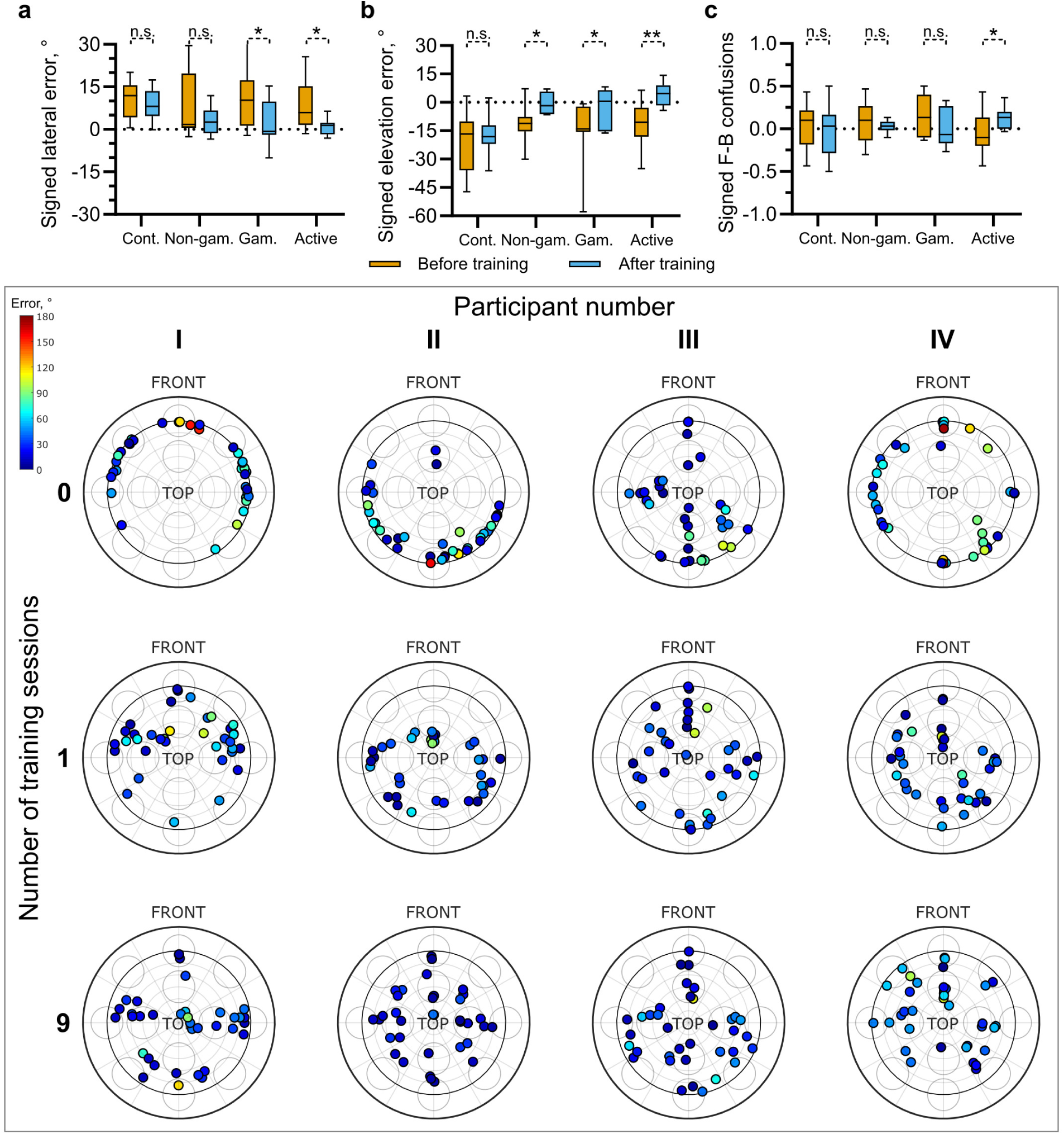
Changes in response biases following training. The top row shows changes in signed lateral (**a**), elevation (**b**) and front-back biases (**c**) both before (orange) and after all nine, 12-minute training sessions (blue). A signed lateral error below zero indicates a tendency to perceive targets more medially and above zero indicates a tendency to perceive targets more laterally. A positive elevation error indicates that targets were perceived above the target position and negative values indicate that they were perceived below. Positive signed front-back (F-B) confusions indicate a tendency to localize targets in rear hemisphere towards the front and a negative value indicates a tendency to localize targets in the front hemisphere to the rear. Indicators of significance are from separate, paired t-tests. **Lower panel**. Distribution of response locations for four example participants before training (top row), following a single, 12-minute training block (second row) and following all nine training blocks (bottom row). Each dot indicates the orientation of a response with 0° azimuth corresponding to straight up on the axes. Elevation is indicated by the distance from the centre, with the origin corresponding to directly upwards and the inner dark ring indicating 0°elevation. The magnitude of the spherical angle error for each response is indicated by the colour of each dot (see colourbar).

One common feature of these measures of response bias is the large degree of variability within each group. This variability is likely to arise from idosyncratic differences between each participants own, natural HRTFs and the non-individualized HRTFs used in the spatialization engine. The lower panel of Fig. 3 shows the locations of all responses given by four example participants, each of whom underwent training, during the first, second and final testing blocks. These data indicate clear response biases during the initial testing block (top row). They also illustrate the high degree of variability in response biases. For example, participant I shows a bias towards the front, giving responses in the rear hemisphere only rarely, while participant II shows the opposite pattern. The responses of participant III lay predominantly close to the sagittal plane, while those of participant IV were well distributed across target azimuth but constrained to low elevation angles. The distribution of responses is much more even following only a single, 12-minute training block (middle row). These clear biases are no longer apparent by the end of training (bottom row).

### Timescale of learning

To investigate the timescale of changes in virtual sound localization accuracy, with a particular focus on any short-term learning effects, testing blocks were carried out between each of the three, 12-minute training blocks on the first day, and following each session of three training blocks on days two and three. The localization errors for each participant group at each of these time points are shown in Fig. 4a. The change in localization error as a function of the number of completed training blocks was fitted with an exponential function of the form *a*exp(−*bx*) + *c* for each group, where *x* corresponds to the number of completed training blocks and *a, b* and *c* were parameters determined using the MATLAB function fit. In this form, the parameters *a* and *c* related to the initial and final localization error and *b* relates to the rate of learning. The fitted functions are indicated by the solid lines in 4a. Adjusted *R*^2^ values indicated that the data were well described by this function for each of the groups undergoing training, 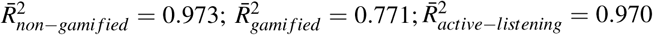. An exponential function provided a poor description of the data for the control group,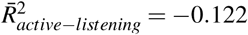. The optimal parameters were found to be *a* = 17.3, 15.7 and 20.7; *b* = 0.270, 0.320 and 0.710; *c* = 23.6, 32.5 and 20.4 for the non-gamified, gamified and active-gamified groups respectively. Taking average values of these fits corresponds to an initial error of 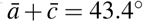, a final error of 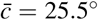, and suggests that errors will reduce by half following 1.60 training sessions (equal to ln(2)*/b*), equating to approximately 19 minutes.

**Figure 4.**
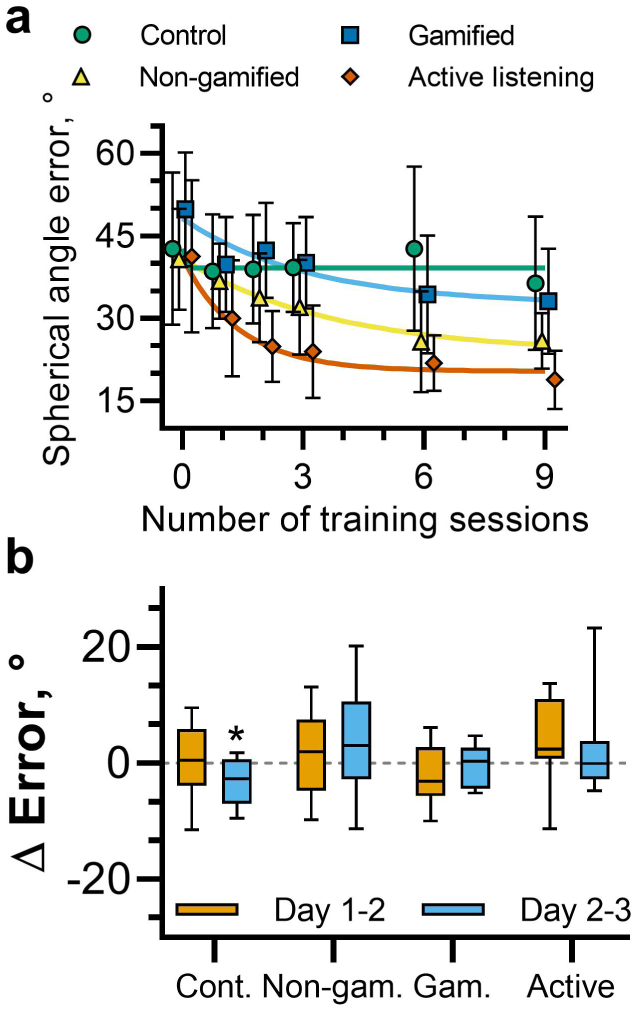
(**a**) Change in spherical angle error as a function of the number of completed 12-minute training blocks. The median spherical angle error was calculated for each participant for each testing block. Shown here are the average values across participants for each group. Error bars indicate the standard deviation. The solid lines represent fitted exponential functions for each group (colours are matched to the symbols). (**b**) Change in spherical angle error between testing blocks from the final testing block on day 1 to the initial testing block on day two (orange) and from the final testing block on day 2 to the initial testing block on day three (blue).

Testing was additionally carried out at the beginning of days two and three, following a gap of up to two days after the previous training and testing session. This facilitated examination of any consolidation or latent learning. These data are shown in Fig. 4b, which illustrates the change in localization error from the last testing block on days one and two and the initial testing block on days two and three respectively, separated by participant group. Separate t-tests were carried out on the change in localization error from days one to two and days two to three. No tests indicated that changes were statistically significantly different from 0°, except for days 2-3 for participants in the control group, whereby localization errors reduced by 3.14° on average, *p* = 0.045. These results demonstrate no evidence of latent learning between sessions for participants undergoing training.

### Generalizability to an untrained HRTF

The previous sections have shown that each of the virtual sound localization training paradigms led to a reduction in localization errors for sounds spatialized using a set of non-individualized HRTFs. To explore whether this learning effect was HRTF-specific, target sounds were also spatialized using a second set of HRTFs within each testing block. The changes in spherical, lateral and polar angle errors, along the change in rates of front-back confusions for sounds spatialized using both the trained and non-trained HRTF sets for each participant group are shown in Fig. 5a-d. Interestingly, the localization error for sounds spatialized using the non-trained HRTFs also decreased in general, and did so such that there was no statistically significant difference between improvements for the trained HRTFs and for the non-trained HRTFs, as determined using separate, paired t-tests.

**Figure 5.**
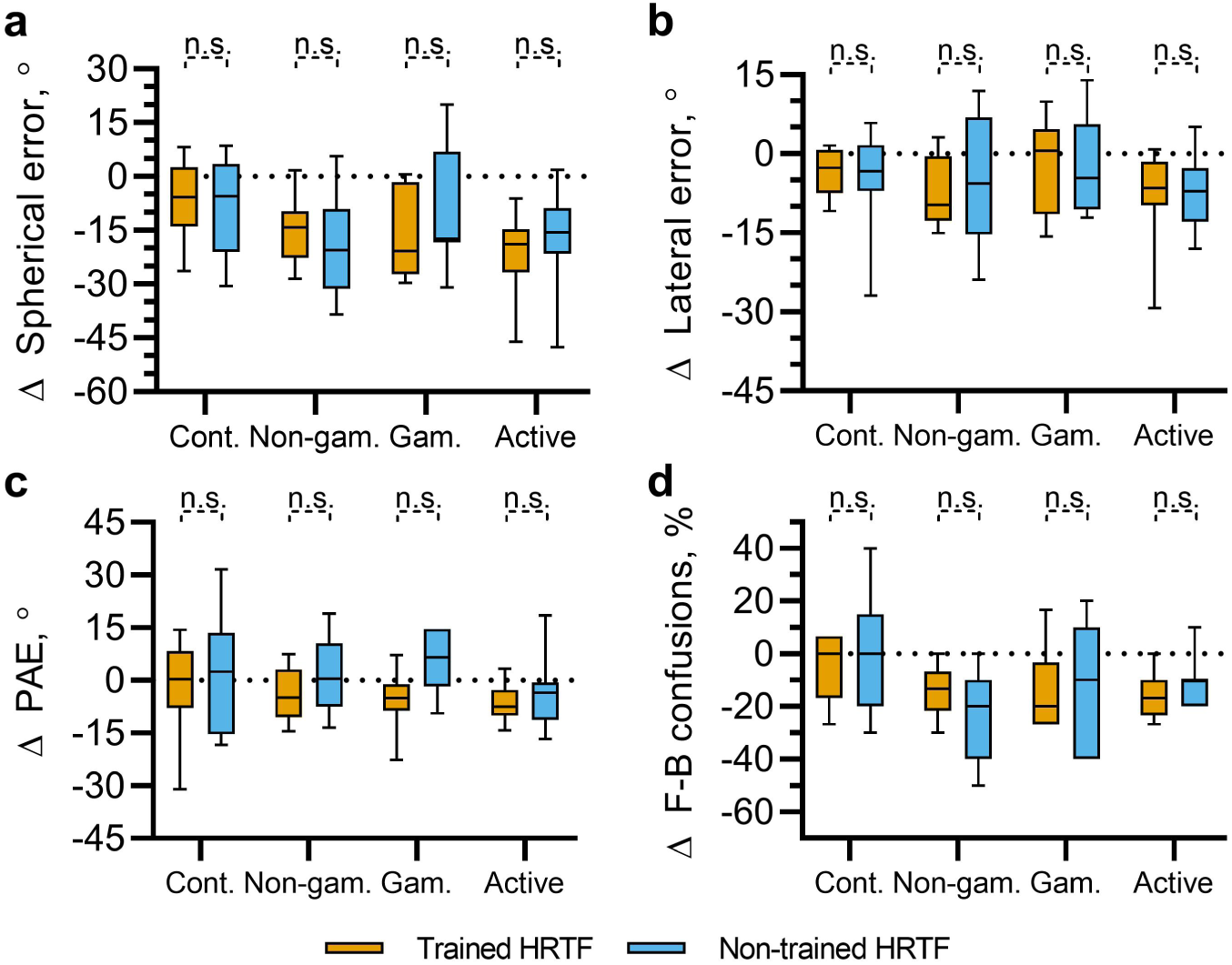
Overall change in spherical angle error, (**a**), lateral error, (**b**), polar angle error (PAE; **c**) and the rate of front-back confusions, (**d**) from the initial to the final testing block for sounds spatialized using the same HRTFs used throughout training blocks (trained HRTF, orange) and for sound spatialized using a second set of non-individualized HRTFs (non-trained HRTF, blue).

We hypothesized that there may be a small, HRTF-specific effect as a result of training that the above analyses were unable to detect. In order to test this, data were pooled across all three training groups. A summary of these data is shown in Fig. 6. Separate paired t-tests were carried out on each measure of localization error (spherical angle error, lateral angle error, PAE and front-back confusion rate). These indicated a significant difference only for the PAE. This difference reflected a larger decrease in PAE for the trained HRTFs (Δ*µ* = −6.74°) relative to the non-trained HRTFs (Δ*µ* = −2.9°).

**Figure 6.**
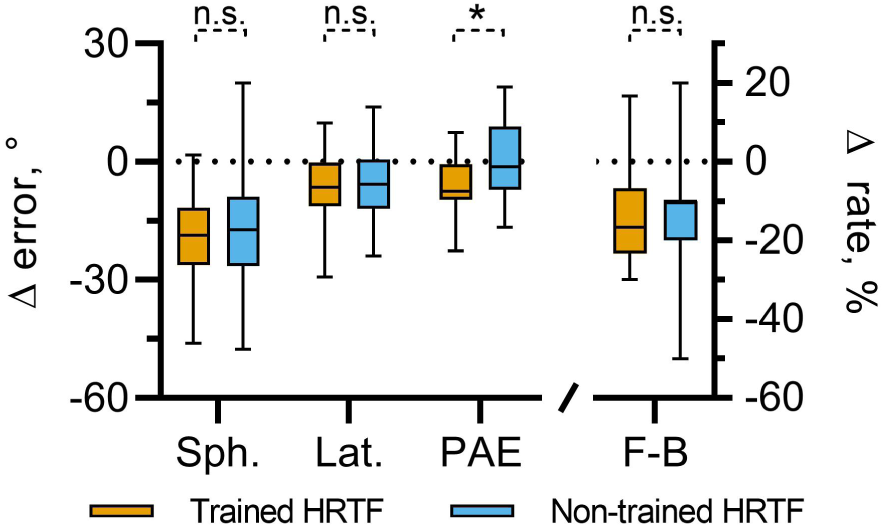
Overall changes in spherical, lateral and polar errors (PAE), and front-back confusion rates (F-B) for sounds spatialized using the same HRTFs used throughout training blocks (trained HRTF, orange) and for sound spatialized using a second set of non-individualized HRTFs (non-trained HRTF, blue). Data are pooled across all participants undergoing gamified, non-gamified and active-gamified training. Significance indicators show results from separate, paired t-tests between the data for the trained and non-trained HRTFs.

It is possible that there were differences in initial localization performance between the two HRTF sets for a given participant. In order to account for these, a one-way ANCOVA was conducted whereby HRTF set was a fixed factor and localization error in the final evaluation session was the dependent variable. Localization error during the initial evaluation session was incorporated into the model as a covariate. This was done separately for spherical angle error, lateral angle error, PAE and front-back confusion rate. In all cases, there was no significant effect HRTFs (trained vs non-trained); Spherical: *F*(1, 51), *p* = 0.410, Lateral: *F*(1, 51), *p* = 0.348, PAE: *F*(1, 51), *p* = 0.694, F-B: *F*(1, 51), *p* = 0.902.

In summary, these data show that the training effect tended to generalize to a second set of non-individualized HRTFs, for which no positional feedback was given, regardless of the measure used. None of the trained groups showed evidence of HRTF-specific effects when considered individually. Pooling across all participants undergoing training provided some evidence of an HRTF-specific learning effect in terms of the PAE. However, after accounting for differences in PAE during the initial testing block, this difference is not statistically significant.

## Discussion

It is now well established that the adult human brain can adapt to altered cues for sound direction. This has been demonstrated in studies using ear plugs^36, 38^ as well as molds inserted in the pinnae^16–18^. Some recent studies have also demonstrated a learning effect using visual positional feedback delivered via a head-mounted display. Zahorik *et al.*^20^ used a very similar training paradigm to those described here and found that the majority of listeners demonstrated improvements in front-back judgements following two, 30-minute training sessions. However, they reported no significant changes in lateral or elevation judgements. Conversely, Majdak *et al*.^21^ reported more general improvements in virtual sound localization precision following training with visual positional feedback, this time using individualized HRTFs. Similarly, Parseihian and Katz^23^ noted improvements in lateral and elevation judgements. Interestingly, this was only the case for the group of listeners using “poorly matched” HRTFs (but not in those using individualized, or “well matched” HRTFs).

The task of quantitatively comparing across studies is non-trivial for several reasons. Firstly, a ubiquitous finding seems to be the large degree of variation across listeners both in localization ability before training and in the effectiveness of training, even when initial ability is taken into account^19^. This makes it likely that findings will be inconsistent, especially where small sample numbers are used. Secondly, differences in pointing method and the distribution and nature of the target stimuli are all likely to have an effect on localization accuracy. For example, here our findings are limited to target sounds primarily in the upper hemisphere with a minimum elevation of 20°below the listener’s interaural axis, whereas the HRTF database includes elevations down to 45°below. Nonetheless, the localization errors before training in this study are consistent with published results where direct comparison is tractable. For example, the average lateral error across all participants before training was 15.1°. This is very close (within ±1°) to those reported by Zahorik *et al*.^20^, and Majdak *et al*.,^39^, who both made use of head-mounted displays and virtual environments. The mean initial front-back confusion rate (40.7%) was also consistent with other studies using non-individualized HRTFs^9, 20^. In summary, these comparisons suggest that the participants recruited here were unremarkable in terms of naive ability.

To facilitate direct quantitative comparisons to previous studies on the effect of training, the various measures of localization accuracy relative to baseline for each participant, separated by group, are shown in Fig. 7. Carlile et al.^18^ perturbed cues for sound localization by inserting molds into the pinnae of human participants, which remained in place continuously during waking hours for 10 days. During this time, some participants underwent training sessions using real sound sources (from a loudspeaker array) and audio-visual positional feedback. Here, we use an identical measure of PAE. Over the course of 10 days, they report a maximum improvement of approximately 5° PAE across all participant groups. We observed a very similar change in PAE for both the gamified and active-gamified groups (Fig. 7c). We also observed somewhat larger changes in rates of front-back confusions; approximately 15% on average across the trained groups compared to a maximum of 10% observed by Carlile et al.^18^. However, it should be noted that the baseline rates of front-back confusion were much lower in their study (40% here compared with approximately 10% on average).

**Figure 7.**
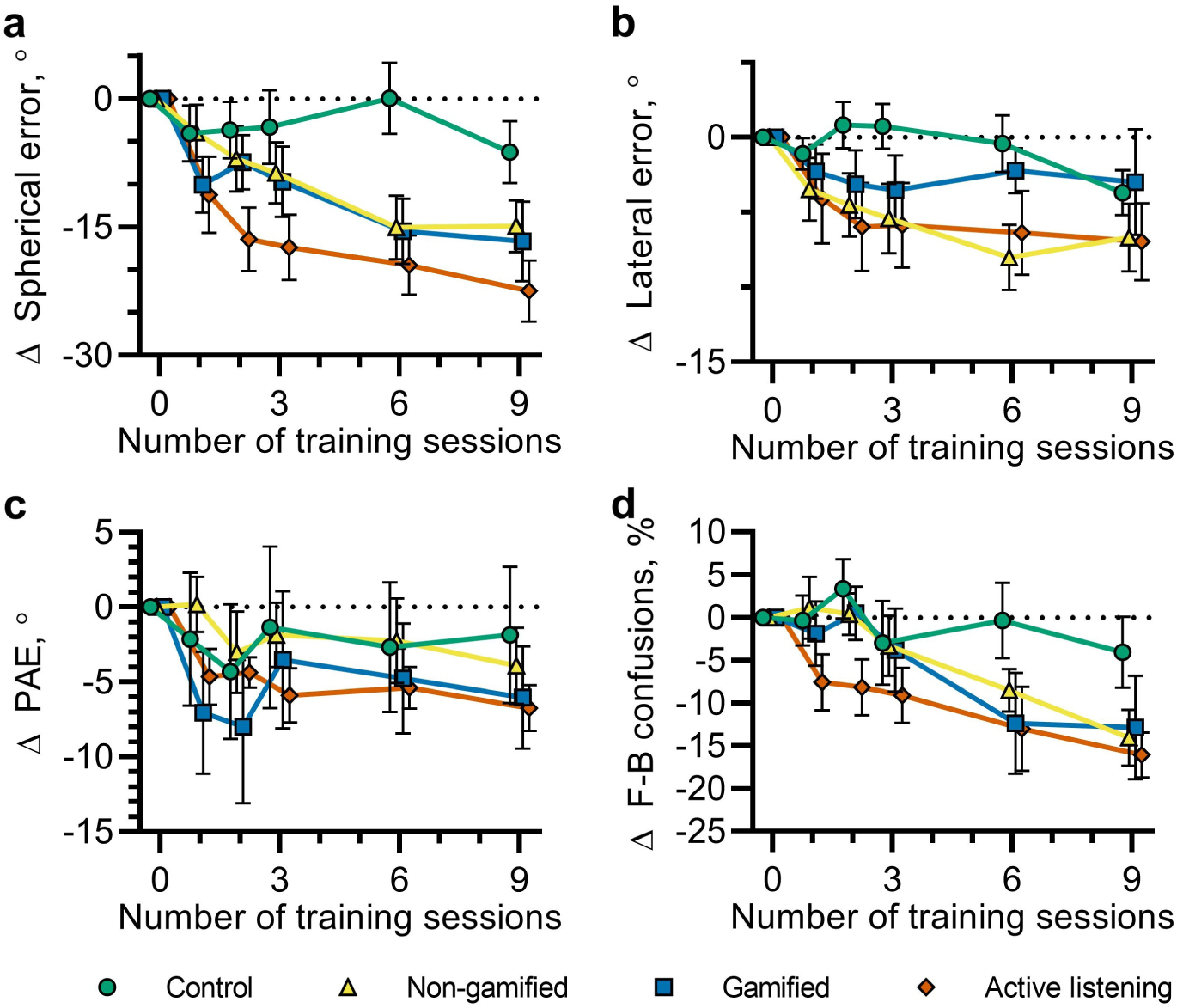
Change in spherical (**a**), lateral (**b**) and polar errors (**c**) and rate of front-back confusions (**d**) relative to baseline for each participant. For spherical, lateral and polar errors, the median error within each testing block was calculated for each participant. Symbols mark the mean of these errors across participants within each group. Error bars indicate standard deviations.

There are some major differences between the present study and that described above. Most notably, they used real, rather than virtual, sound sources and were interested in changes over a considerably longer timescale. Stitt et al.^37^ employed a more similar approach, whereby the ability of participants to localize virtual sounds, spatialized using non-individualized HRTFs, was trained and tested. Their study also included a group using individualized HRTFs. All participants underwent localization training. Despite participants undergoing training of a similar total duration (10 sessions of 12 minutes), Stitt et al.^37^ report larger changes on average in PAE than observed here or in the study described above^18^. One participant group showed a 15° improvement on PAE, using an identical measure. It should be noted, however, that this is in the context of high polar errors before training, particularly for listeners using non-individualized HRTFs (although a direct comparison is not possible since this study presents a different measure of polar error for absolute values to that used to describe changes relative to baseline^37^). All participant groups in that study showed little to no improvement in lateral error, whereas the non-gamified and active-gamified participants in the present study showed an improvement of over 5° (Fig. 7b). In general, such differences between studies are unsurprising given the high degree of variability between subjects both in terms of absolute localization accuracy, particularly at baseline, and in terms of improvement.

This study aimed to explore the potential of “gamification” to enhance the effects of training on adaptation to non-individualized HRTFs. It has previously been shown that videogame-like paradigms can be used to effect improvements in virtual sound localization^31^. Here, we compared a standard, adaptive psychoacoustic training paradigm to “gamified” version of the same task, which incorporated performance-related feedback by, amongst other things, awarding points for positive actions (“hits”) and decreasing player “health” for negative actions (“misses”). However, the introduction of these elements alone was insufficient to effect a significantly better outcome than the more traditional version. It has been previously shown that game-play can have an enhancing effect on perceptual learning, even where the video game is not directly related to the learning task, or indeed involving the same modality^34, 40^. However, it may be that our “gamified” version was not sufficiently distinct from the standard task, since both versions provided performance-related feedback in form of “hits” and “misses” and, anecdotally, participants reported being quite motivated to improve over time whether they were using the gamified version or not. Since perceptual learning enhancement appears to be effected through videogame play regardless of relation to the learning task, it is possible that an alternative approach might be to incorporate existing games into the localization training regimen, or to design games primarily for fun rather than designing the game around the localization task. A further consideration is that there may be substantial variability in how well individuals respond to gamification. Anecdotally, few of our participants indicated a strong interest in gaming and thus it is possible that this group did not respond well to the reward mechanisms typically utilised in game design. Future studies could recruit distinct cohorts of “gamers” vs “non-gamers” to test the idea that gamers could be a self-selecting group who respond well to gamification in a perceptual learning paradigm.

Active listening, on the other hand, does seem to robustly lead to a more marked improvement. Here, active listening refers to listener’s opportunity to experience a stationary virtual sound source while being free to move their head. This necessitates the use of head tracking, whether that is through a dedicated head tracking system, or simply using positional data from embedded sensors in a smartphone, as used in the present study. The data presented here suggest that the experience of ongoing, relative motion may have an enhancing effect on virtual sound localization training. It should be noted that the active listening group were presented with the stimuli repeatedly during a trial so it could be argued that this is simply an effect of greater stimulus exposure. However, we found no significant relationship between the total number of times each participant was presented with spatialized stimuli during training and the overall improvement in localization accuracy. This suggests the time engaged with the training task seems to be more important than the absolute number of trials completed.

This study was designed such that it was possible to measure any putative consolidation or latent learning effects. Since localization testing took place at the beginning and end of each training day, it was possible to measure any change in virtual sound localization ability occurring during rest periods, during which no training took place. A consolidation effect was observed, whereby improvements were maintained between sessions. Although latent learning (improvement on a task occuring during rest periods) has been observed in a previous study on auditory perceptual learning, albeit on a simpler task^41^, we observed no evidence for it here. Molloy *et al.*^41^ reported that very short training sessions (∼8 minutes) can optimise latent learning and can therefore be the most efficient regimen. It is possible that shorter training sessions spread over more days might, indeed, be optimal and could be appropriate in some virtual audio use cases.

Along with several other studies, we have demonstrated that training leads to improvements in several measures of virtual sound localization accuracy. For some applications (e.g. gaming or virtual auditory display), the mechanisms by which these improvements occur are not important. However, if virtual audio is to have a clinical application, it would be important to have a deeper understanding of these. This could also have implications for virtual audio system design. Here, we consider the putative contributions of two types of learning process; procedural and perceptual. By procedural learning, we refer to improvements related to the “response demands” of the task^42, 43^. In this case, the response demands comprise a perceptual-to-motor mapping required to accurately turn and orient towards the perceived direction of a sound source. Perceptual learning refers to a number of putative mechanisms associated with an auditory-to-spatial mapping. Here, this involves learning to associate unfamiliar acoustic cues with specific sound source directions. In virtual sound localization tasks, this is often referred to as “HRTF adaptation”. Firstly, we will consider the evidence for and against the interpretation of our data as indicative of procedural learning. We will then discuss experiment design considerations for disentangling these effects in future.

In comparison to many previous studies on adapting to altered cues for sound localization, this study focused on changes over a very short timescale. Typically, such adaptation has been measured over the course of hours, days and weeks (for a review see Mendonça *et al.*^19^). In agreement with the few studies that have also focused on short-term changes, we show that significant changes occur after a total training duration of the order of an hour spaced over three days^20, 22^–^24, 37^. This is an encouraging result if future systems using generic, non-individualized HRTFs are to find broader application, since it seems likely only a minority of use cases will justify longer adaptation periods. However, such rapid changes are often attributed to procedural learning^42^. Furthermore, we show that the majority of our learning effect generalised to a second, non-trained HRTF. This is also consistent with a predominantly procedural (i.e. perceptual-motor) mapping. Furthermore, Honda *et al.* (2007)^31^ trained listeners using a virtual sound localization game and found that this resulted in improved free-field localization. They went on to interpret this as evidence of a perceptual-motor (procedural) learning effect.

Hawkey *et al.*^43^ explicitly explored the question of whether perceptual learning effects can occur on very short timescales. They reported that auditory perceptual learning on a frequency discrimination task did indeed occur rapidly (within 500-600 trials; comparable to approximately two training sessions in the present study). Furthermore, Trapeau *et el.*^44^ trained participants in a free-field localization study using a spherical loudspeaker array, whereby one group of participants had their spectral cues altered using ear molds and another underwent the same training and testing procedure without ear molds. After six, 15-minute training sessions over six days, they observed improvements in elevation and lateral angle judgements in the participants with ear molds comparable to those reported here. They observed no learning effect for the participants without ear molds, suggesting that the procedural learning effect in a comparable sound localization task, whereby responses are given by head-pointing, was minimal. One reason for this could be that oreinting towards the perceived direction of a sound is a fairly natural action, which is even displayed by newborn human infants^45^ (as opposed to the swatting action employed by Honda *et al.*^31^).

In summary, it seems reasonable to posit that at least some of the learning effects observed here are due to auditory perceptual learning, despite the observation that it generalizes to both the trained and non-trained HRTFs. However, it is worth considering how one could be certain that observed changes in virtual localization accuracy following training are indeed associated with “HRTF adaptation”. One approach is to include a group of listeners who are presented with sounds using their own, individualized HRTFs. One might then assume that only the differences between this and a second group using non-individualized HRTFs are due to HRTF adaptation. For this assumption to hold true, it is necessary that there is no perceptual difference between a listener’s “true” HRTFs and the measured, individualized ones. However, it is not clear that this is the case, since listeners using individualized HRTFs do show some localization deficits before training; poorer elevation judgements and higher rates of front-back confusions^1, 8^. The approach used by Trapaeu *et al.*, whereby training effects for participants trained using modified spectral cues are compared with those for participants using unmodified cues in the free field, avoids this issue and doing so yields no evidence of a significant procedural learning effect for a sound localization, head-pointing task. A complimentary approach would be to design a testing procedure that relies on adapting to non-individualized HRTFs, but is distinct from the localization training task, perhaps using a different method of indicating responses, a distinct class of stimuli or a different task altogether. One possibility is to use a task that relies of a different aspect of spatial processing (e.g. spatialized speech recognition with spatialized masking sounds), although this would rely on relative rather than absolute cues for sound location so it is unclear how effective an absolute localization training regimen would be.

Previous studies have shown that localization training paradigms can lead to learning effects that generalize to novel stimulus-position pairings^24^, and that training on a fitted HRTF can lead to an improvement in free field sound localization^31^, but to our knowledge generalization across unfitted HRTFs has not been investigated directly. One reason one might expect the observed generalization is that the learning effects are primarily due to procedural learning, but for the reasons outlined above, it is reasonable to expect that this does not account for all of the improvements observed. This effect might also be observed if the HRTF sets used in this study happened to be perceptually similar. The method used to select them from the HRTF database would certainly not guarantee perceptual distinctiveness; the subset they were taken from was selected on a somewhat utilitarian basis whereby they tended to produce the most subjectively realistic, spatialized percepts for the greatest number of listeners^14^. It could be argued that this method would tend to produce subsets that epitomize stereotypical features and minimize idiosyncratic variation. Indeed the two chosen HRTFs appear to be relatively close in terms of perceptual similarity for median plane trajectories^46^, though it should be noted that a later study from the same team noted that such subjective HRTF ratings were not reliably repeatable, even for expert listeners^47^. To discriminate between changes due to HRTF-specific adaptation and other factors, it may be useful to be able to select HRTFs that are maximally perceptually distinct. This is non-trivial, especially where the cues used for discriminating the specific target orientations used are complex and poorly defined. One approach could could be to use a clustering analyses whereby HRTFs are defined using acoustic dimensions corresponding to putatively perceptually relevant cues for direction, such as that proposed by So et al.^48^, but more work on perceptual similarity of HRTFs would be required before such a procedure could be considered robust.

One possibility that could account for the observation that the adaptation appears to generalize to more than one HRTF is that the process involves reweighting acoustic cues for sound source location. In this scenario, listeners would learn to rely less on idiosyncratic cues specific to their own HRTFs and more on acoustic cues that are shared with the non-individualized ones. An illustrative example might be that listeners begin to rely on interaural level differences more than time differences for lateral angle judgements, if these are indeed perceptually more robust. Such a mechanism relies on redundancy in auditory-spatial cues and has been put forward as a process underlying auditory perceptual learning in other contexts^36, 49^. This idea seems to be supported by the rapid change in response bias observed here. It could be that, during the initial testing block, participants did not receive the cues they would normally rely on for the most accurate localization leading to diverse “guessing” strategies. However, upon receiving some positional feedback in a single training block, they were able to rapidly switch listening modes to place greater emphasis on cues that generalized between their own and the unfamiliar, non-individualized HRTFs. Indeed, such a rapid switch of listening mode can be observed in listeners switching between complex listening environments^50^. Cue reweighting also provides an explanation of the lack of after-effect routinely observed in studies that temporarily alter sound localization cues (using molds, for example). Previous interpretations have been that listeners develop a many-to-one auditory-place mapping, effectively comprising parallel, internal representations of multiple HRTFs for a given sound direction^44^, the ecological benefit of which is unclear. On the other hand, cue reweighting would be an adaptive strategy in the real world for dealing with certain environments (e.g highly echoic ones where binaural cues could be profoundly disrupted) or sounds (e.g. those without energy in the frequency ranges required to make use of spectral notches).

One of the aims of this project was to demonstrate that this adaptation could be achieved using readily available, consumer electronics. Previous studies on adapting to non-individualized cues for virtual sound location have utilised specialist facilities and equipment typically involving dedicated head tracking equipment^23, 39, 44^ or *ad hoc* visual displays^35^. Whilst dedicated platforms may have the potential for greater experimental control and accuracy, more pragmatic solutions will be required if non-individualized virtual audio is to find more widespread application. Here we demonstrate that the accuracy of localization using consumer equipment is comparable to previous studies and it is possible to effect and measure improvements in virtual sound localization accuracy using non-individualized HRTFs in the same way. Since developing the platform used in this study, which utilised both a PC and a smartphone, several systems have become freely available that implement HRTF-based virtual sound spatialization entirely on a single mobile device (e.g.^51^), which make it relatively straightforward to implement a similar platform using only a smartphone and headphones. This simplicity raises the possibility that virtual sound systems could have broader application than previously thought. For example, it is possible to imagine research, or even clinical, applications whereby the effectiveness of hearing prostheses could be tested in controlled, virtual environments without the need to install bulky, expensive loudspeaker arrays. Of course, the effectiveness of such applications is only assured if it is possible to assume that the reduction in localization error observed following brief periods of training corresponds to an adaptation mechanism whereby the cues and strategies used are similar to those used in real-world listening. It follows that one limitation of our approach is the use of localization accuracy as a proxy measure for “HRTF adaptation”. If the goal is to use the brain’s ability to adapt to broaden the application of virtual sound systems either in consumer products, auditory research or clinical applications (such as audiometry in simulated real-world environments), then the quantity we are actually interested in is perceptual “realism”. This is ill-defined and difficult to measure reliably^52^, which is presumably why localization accuracy is commonly used. The extent to which this proxy measure is associated with perceptual realism, however, is not clear.

In summary, we have demonstrated a virtual sound system that uses only readily available consumer electronics. We implemented training programmes that led to significant decreases in virtual sound localization error following fewer than than two hours of training (split over three sessions). We show that active listening, the ability for listeners to move their head relative to virtual sound sources, appears to enhance the effects of this training. The majority of improvements in sound localization accuracy generalized to a second, non-individualized HRTF set, which was not used in training. Only a small proportion of changes in polar angle judgements were HRTF-specific. This may suggest that the observed improvements in virtual sound localization following training do not wholly reflect a mechanism whereby listeners learn a parallel internalised auditory-spatial map. Instead, this observation could be explained by an underlying mechanism that involves switching to a different localization strategy for sounds spatialized using non-individualized HRTFs by cue reweighting. This would imply that listeners showing HRTF adaptation as measured by localization accuracy may not be using a naturalistic listening mode, which has implications for possible future applications of virtual audio technology.

## Methods

### Participants

All procedures were reviewed and approved by the Imperial College Research Ethics Committee and all research was performed in accordance with the relevant guidelines and regulations. At the start of the experiment, participants received an information sheet outlining the experiment and were informed that they may leave the study at any time. After this, each participant provided informed consent to take part in the experiment. Before entering the study, participants were also asked to complete a questionnaire, which revealed no reported cognitive or auditory deficits. All data were anonymized.

A total of 36 participants (14 females and 24 males), aged between 18 and 38, were recruited for this study. Participants were randomly assigned to one of four groups. Three of the groups underwent sound localization testing and training in a virtual environment, each utilizing a different training paradigm. The fourth group acted as a control and remained in a quiet room between testing sessions.

### Stimuli

A set of 19 acoustically complex stimuli were developed. The sounds were designed such that they were consistent with the virtual environment and contained rich cues for localization. As such, they were designed to resemble a short radio communications transmission. The stimuli comprised a combination of pink (1/f) noise, a segment of speech produced by a male talker and a 1 kHz tone. A schematic of the stimulus is shown in Fig. 8a, and an example is provided (Supplementary Audio). An initial 200 ms noise burst was followed by a one second fragment of continuous Italian speech with low level pink noise, another 200 ms noise burst and, finally, a 200 ms, 1 kHz tone. Each stimulus comprised unique noise and speech fragments. Each segment was ramped on and off using a 10 ms raised-cosine ramp. From this set, a single stimulus was used only during testing, whilst the other 18 stimuli were selected from randomly during training.

**Figure 8.**
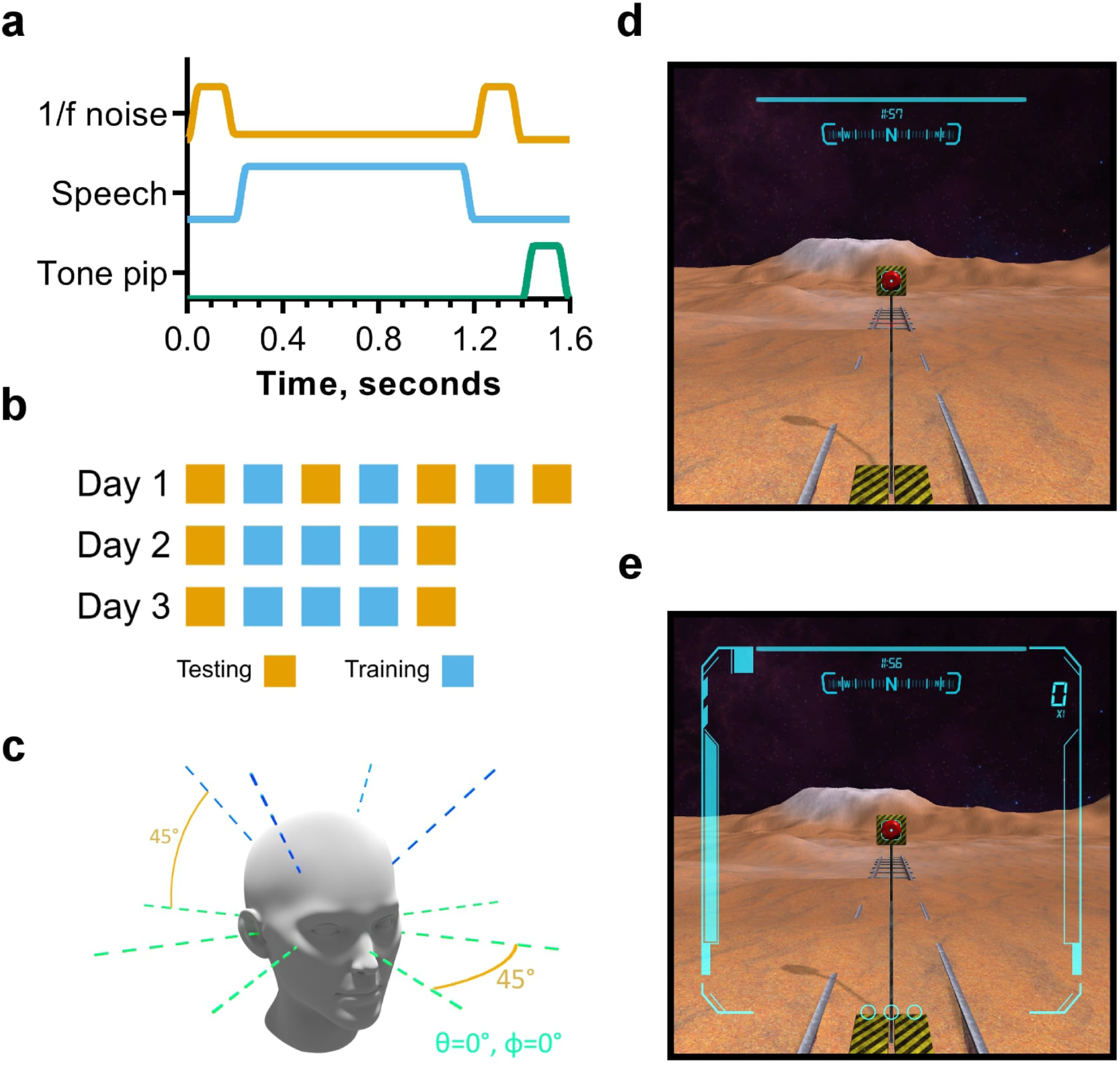
(**a**) Schematic representation of the complex stimulus used during training and testing. **b**) schematic of the experimental design indicating the block structure of each session. Testing blocks are in orange and training blocks in blue. Each session was carried out on a different day. **c** Diagram of the centroids of target orientation “regions” used in testing blocks. Target sounds deviated from these centroids by up to 20°. **d**,**e**) Screenshots of the virtual participants view in the virtual reality application. The marked features correspond to a) timer, b) cardinal direction indicator, c) current score, d) player health indicator, e) animated “charge” indicator visual effect, f) consecutive hit counter. **d** shows the HUD used in the non-gamified version and **e** shows the HUD for the gamified and active-listening versions.

Sounds were spatialized using HRTFs from the IRCAM Listen database^53^. Two HRTFs were randomly selected from a subset of this database, which was determined in a previous study to contain the seven HRTFs that produced the best subjective spatialization for the most listeners^14^. These corresponded to participant numbers IRC0008 and IRC0013 in the database. The former was used to spatialize sound during both training and testing. The latter was used only during testing. The interaural time differences were left unmodified, as per the original HRTFs. All stimuli were generated and stored in 44.1 kHz, 16-bit format.

### Spherical coordinate systems

Throughout this manuscript, two spherical coordinate systems are used. The first is a single-pole system whereby sound source coordinates are specified using two coordinates; azimuth, *θ* and elevation, *ϕ*. In this coordinate system, the azimuth refers to a rotation around a vertical axis passing through the centre of the head with 0° being directly in front and 90° being directly to the left. Elevation is the angle from the horizontal plane with 0° being horizontal and 90° being directly upwards. This coordinate system is intuitive and mathematically convenient and is therefore used in the subsequent description of target orientation, as well as in the experiment software.

A second coordinate system is used in the data analysis, referred to variously in the literature as “head-related”^8^, “double-pole”^54^ or “interaural-polar”^55^. This coordinate system was inspired by binaural hearing and also involves two coordinates; a lateral coordinate that defines the angle between an “interaural axis” (coincident with the line linking the listeners ears) and the sound source, and a polar coordinate that defines the angle formed between the horizontal plane and a line linking the sound source and the interaural axis, perpendicular to that axis. In binaural listening, the dominant cues for the lateral coordinate are considered to be the interaural time and level differences and the dominant cues for the polar coordinate are considered to be spectral since it defines a point on a circular cross section of the “cone of confusion”^56^. See reference^54^ for a diagram illustrating these coordinate systems. In this manuscript, we distinguish the coordinate systems using the nomenclature of “azimuth” and “elevation” for the single-pole system, and “lateral” and “polar” for the double-pole, interaural system.

### Experiment design and procedure

For each participant, the experiment comprised three sessions, each completed on a different day and separated by no more than two days. Each session incorporated sound localization testing and training blocks in a virtual environment. During a session, participants sat on a freely rotating chair in the centre of a dark, quiet room. The virtual environment was presented using a head-mounted display, and auditory stimuli were presented over headphones. Participants interacted with the experiment software using a gamepad. For details of the equipment, see **Equipment and the virtual sound localization environment**. During testing and training blocks, participants initiated trials in their own time by orienting towards a button within the virtual scene and activating it using the gamepad. Doing so initiated playback of a randomly selected complex auditory stimulus (see **Stimuli**).

Testing and training blocks were differentiated by whether positional feedback, in the form of a visual sound source positional indicator, was given. Participants underwent an initial testing block at the start of each session, followed by three 12-minute training blocks, followed by a final testing block. Participants were encouraged to take a 5-10 minute break between blocks, during which they remained in the quiet room. In order to capture any very rapid learning effects, additional testing blocks were carried out between each of the training blocks on the first day. This design is represented schematically in Fig. 8b. The control group followed the same process, but remained in the quiet testing room during the periods in which the other groups underwent training, during which time they were allowed to engage in non-auditory activities such as reading.

In training blocks, stimuli were spatialized using randomly generated source locations uniformly distributed over the upper hemisphere by setting *θ* = 2*πu* and *ϕ* = *sin*^−1^*v*, where *θ* and *ϕ* are the azimuth and elevation angles respectively and u and v are random variates uniformly distributed on the interval [0, 1]. The choice of spatializing sources only in the upper hemisphere was made because the virtual (and indeed real) environment had a floor, and we wanted to avoid providing conflicting audio-visual cues. The participants were instructed to remain oriented in the same direction throughout the 1.6 seconds of stimulus playback. If the orientation of the head-mounted display deviated by more than 2.5° in any direction during this time, the trial was cancelled and the training did not continue until the participant initiated a new trial (with a new source location). Following stimulus playback, participants were instructed to orient towards the perceived direction of the sound source and indicate their response using a button on the gamepad. When the response was given, the stimulus was played back a second time, spatialized at the same source location.

Visual positional feedback was presented simultaneously by introducing a spherical object in the virtual scene at the target sound source location after each participant response. If the response was within the bounds of the object, the response was indicated to the participant as a “hit”, otherwise it was a “miss”. The way this was visualized depended on the training paradigm used and is detailed in the following section. The size of the target object varied adaptively. The initial target size was set such that responses were indicated as a “hit” if there was less than 25° deviation from the target in any direction. After achieving three consecutive hits, the target size decreased by 10%. After five misses at a given target size, the target size reverted to the previous one until reaching the initial size. The radius of the target object was therefore given by *r* = 0.9^*L*−1^*dsinθ*, where *L* is the current difficulty level, *d* is the target distance and *θ* is the allowed angle of deviation for a “hit”. This mechanism of “hits” and “misses” was used primarily to provide some performance-related feedback - in all subsequent analyses only the difference between the response and target orientation was used as the variable of interest. All participant groups undergoing training were instructed to initiate and complete trials continuously for 12 minutes per training block.

Testing blocks were carried out using the same virtual environment and process, except positional feedback was not provided following participant responses. Further, testing blocks comprised a fixed number of trials rather than a fixed time limit. In order to ensure consistency across participants, target stimuli were positioned systematically. Eight orientations were defined at 0° elevation, with azimuths were separated by 45° increments beginning at 0°. A further four orientations were defined at 45° elevation with azimuths equal to 45°, 135°, 225° and 315°. These orientations are visualized in Fig. 8c. In order to avoid direct stimulus to response mapping, targets randomly deviated from these orientations by up to 20°. In one testing block, four targets were presented for each target orientation with a different, random deviation each time. Three of these were spatialized using the same HRTFs used in the training blocks. In order to test if learning effects transferred to more than one set of HRTFs, the fourth target was spatialized using a second HRTF set for which the participants had received no positional feedback.

### Training paradigms

Three versions of the training software were developed. Each utilised an identical virtual environment. The first, basic version, referred to as “non-gamified”, presented participants with a simple head-up display (HUD) incorporating a compass to show absolute orientation as a cardinal direction and a timer, showing the time remaining (Fig. 8d). In this version, visual positional feedback was implemented using plain, spherical objects of uniform colour (green for a “hit” and red for a “miss”). If the target was not in the visual field of the participant when they gave their response, an arrow on the HUD indicated the direction to the target following the shortest path. A second “gamified” version was also developed, which incorporated several videogame design elements. These comprised a score indicator, a consecutive hit counter and player “hit points” (Fig. 7e). Points were rewarded for target hits. When the target size decreased following the adaptive procedure described above, this was indicated as a level progression to the participant using a sound and text on the HUD (i.e. “LEVEL 2”). In this version, the positional feedback was provided by an animated spherical “robot” in the visual scene, which also fired a “laser” at the player if a response was a miss, leading to a decrease in player “health”. The laser had an accompanying sound effect, which was not spatialized. When the health ran out, this corresponded to an increase in target size, as per the adaptive procedure. A “game over” overlay was presented to the participant and the “hit points” were reset before immediately continuing with training.

In both of these training paradigms, the participant/player was required to keep their head oriented in the same direction during target stimulus playback (within 2.5° deviation) otherwise the trial was cancelled. However, there is some evidence to suggest that “active listening” - the ability of the listener to move their head relative to a sound source - plays an important role in adapting to modified cues for sound localization^19^. A third version of the training software was implemented that incorporated this. There was no requirement for participants to remain oriented in the same direction during stimulus playback, which was looped until listeners gave their response. Participants were not explicitly encouraged to move their head to better localize sound sources. This “active listening” version was in all other ways identical to the gamified version described above. Note that all participants utilised head tracking and were able to move their head relative to the sound source when receiving positional feedback upon giving a response. All participants used exactly the same version of the testing software. A video of these training paradigms as well as the testing paradigm is available online (Supplementary Video). Please note that due to the screen capture software and file size limitations, the audio in the video is not representative of the audio generated by the spatialization software.

### Equipment and the virtual sound localization environment

The virtual environment was rendered stereoscopically on a smartphone-based (iPhone 6s) head-mounted display. Participants interacted with the phone using a handheld controller connected via Bluetooth (Mad Catz C.T.R.L.i). Head-tracking data was transmitted via wireless Ethernet connection (using OSC - Open Sound Control) to a PC that handled spatial audio rendering. Sound playback and real-time binaural spatialization were implemented using the LIMSI Spatialization Engine^57^ - a real-time binaural audio spatialization platform based on Cycling74’s Max/MSP. Binaural audio was presented via a Focusrite Scarlett 2i2 USB audio interface using Sony MDR 7506 closed-back headphones. The head-tracking data was internally refreshed at a rate of 2000Hz, and the frame size for the audio spatialization was set at 128 samples. The delay caused by the wireless connection between the head-tracker and the PC was variable (±2 ms), but the overall latency of the whole system was never above 20 ms. The inertial sensors and magnetometer on the iPhone 6s allowed for a maximum tracking error of ±1°.

A virtual, moon-like environment was designed to be acoustically neutral to minimize the potential mismatch between the anechoic stimuli and the perceived acoustic properties of the virtual space. The scene was also populated with some landmarks as it has been shown that a lack of visual frame of reference is detrimental to sound localization accuracy^58^. Screenshots of part of the environment are shown in Fig. 7d and Fig. 7e, which show the non-gamified and gamified HUD respectively. The virtual environment can be seen in more detail in the Supplementary Video.

### Measures of localization error

Four measures of localization error were used. These are (1) the angle formed by a vector oriented towards the location of a target, virtual sound source and a vector oriented in the direction of the listeners head when indicating their response. This is referred to as the spherical angle error. (2) the absolute difference between the target lateral angle and response lateral angle, both in the interaural coordinate system. This is referred to as the lateral error. (3) The polar angle error (PAE), which is the same as that described by Carlile *et al.*^18^. This is equal to the difference in polar angle between the target and response in the interaural coordinate system, which were only calculated for responses that were not front-back confused and were scaled by the cosine of the target lateral angle (to account for changes in the sizes of cones of confusion). (4) The front-back confusion rate, which was given by the percentage of responses in the incorrect hemisphere (front vs rear). Trials where the target had an absolute lateral angle >65° (within 25° of the interaural axis) were not included in this calculation. Again, this metric is the same as that used by Carlile *et al.*^18^.

## Acknowledgements

This work was supported by the 3D Tune-In project^59^, part of the Horizon 2020 research and innovation programme under grant agreement No. 644051. The authors would like to thank Brian F.G. Katz and his team at LIMSI-CNRS for the use of the LIMSI Spatialization Engine.

## Author contributions statement

M.A.S, L.P & D.F.M.G developed the study concept and experiment design. M.A.S. designed and build the experimental setup including instrumentation and the virtual environment software. Data was collected by M.A.S., J.H.L & C.H. and analyzed by M.A.S. with input from C.K. Analysis scripts were produced by M.A.S. with input from C.K. M.A.S. produced the manuscript and figures with input from L.P. and D.F.M.G.

## Data availability

The Unity project used to build the software for the iPhone 6 and the full dataset, including Matlab scripts to read it, are publicly available on Zenodo^60^. Please note that the authors do not have permission to redistribute the LIMSI Spatialization Engine, which is required to reproduce the stimuli. Please contact the authors for details.

